# Feasibility of Automated Deep Learning Design for Medical Image Classification by Healthcare Professionals with Limited Coding Experience

**DOI:** 10.1101/650366

**Authors:** Livia Faes, Siegfried K. Wagner, Dun Jack Fu, Xiaoxuan Liu, Edward Korot, Joseph R. Ledsam, Trevor Back, Reena Chopra, Nikolas Pontikos, Christoph Kern, Gabriella Moraes, Martin K. Schmid, Dawn Sim, Konstantinos Balaskas, Lucas M. Bachmann, Alastair K. Denniston, Pearse A. Keane

**Author notes:** Shared first authorship (Livia Faes, Siegfried K. Wagner). **Corresponding author:** Pearse A. Keane, MD FRCOphth, NIHR Biomedical Research Centre for Ophthalmology, Moorfields Eye Hospital NHS Foundation Trust and UCL Institute of Ophthalmology, United Kingdom.

## Abstract

Deep learning has huge potential to transform healthcare. However, significant expertise is required to train such models and this is a significant blocker for their translation into clinical practice. In this study, we therefore sought to evaluate the use of automated deep learning software to develop medical image diagnostic classifiers by healthcare professionals with limited coding – and no deep learning – expertise.

We used five publicly available open-source datasets: (i) retinal fundus images (MESSIDOR); (ii) optical coherence tomography (OCT) images (Guangzhou Medical University/Shiley Eye Institute, Version 3); (iii) images of skin lesions (Human against Machine (HAM)10000) and (iv) both paediatric and adult chest X-ray (CXR) images (Guangzhou Medical University/Shiley Eye Institute, Version 3 and the National Institute of Health (NIH)14 dataset respectively) to separately feed into a neural architecture search framework that automatically developed a deep learning architecture to classify common diseases. Sensitivity (recall), specificity and positive predictive value (precision) were used to evaluate the diagnostic properties of the models. The discriminative performance was assessed using the area under the precision recall curve (AUPRC). In the case of the deep learning model developed on a subset of the HAM10000 dataset, we performed external validation using the Edinburgh Dermofit Library dataset.

Diagnostic properties and discriminative performance from internal validations were high in the binary classification tasks (range: sensitivity of 73.3-97.0%, specificity of 67-100% and AUPRC of 0.87-1). In the multiple classification tasks, the diagnostic properties ranged from 38-100% for sensitivity and 67-100% for specificity. The discriminative performance in terms of AUPRC ranged from 0.57 to 1 in the five automated deep learning models. In an external validation using the Edinburgh Dermofit Library dataset, the automated deep learning model showed an AUPRC of 0.47, with a sensitivity of 49% and a positive predictive value of 52%. The quality of the open-access datasets used in this study (including the lack of information about patient flow and demographics) and the absence of measurement for precision, such as confidence intervals, constituted the major limitation of this study.

All models, except for the automated deep learning model trained on the multi-label classification task of the NIH CXR14 dataset, showed comparable discriminative performance and diagnostic properties to state-of-the-art performing deep learning algorithms. The performance in the external validation study was low. The availability of automated deep learning may become a cornerstone for the democratization of sophisticated algorithmic modelling in healthcare as it allows the derivation of classification models without requiring a deep understanding of the mathematical, statistical and programming principles. Future studies should compare several application programming interfaces on thoroughly curated datasets.

## INTRODUCTION

Diagnosis depends on data: its collection, integration and interpretation enables accurate classification of clinical presentations into an accepted diagnostic category. Human diagnosticians achieve acceptable accuracy in such classification tasks through the learning of diagnostic rules (patterns recorded by other human diagnosticians) followed by training on real cases for which the diagnostic labels are provided (supervised clinical experience). Due to the limited capacity of human neural networks (brain), the amount of data utilized to create these diagnostic rules, and then to reach a diagnosis on an individual patient is highly selective and biased, with the vast majority of available data being ignored. In artificial intelligence (AI), the technique of deep learning uses artificial neural networks – so-called because of their superficial resemblance to biological neural networks – as a computational model to discover intricate structure and patterns in large, high dimensional datasets such as medical images.[1] A key feature of these networks is their ability to fine-tune based on experience, allowing them to adapt to their inputs, thus becoming capable of learning. It is this characteristic which makes them powerful tools for pattern recognition, classification, and prediction. In addition, the features discovered are not predetermined by human engineers, but rather by the patterns they have learned from input data.[2] Although first espoused in the 1980s, deep learning has come to prominence in recent years, driven in large part by the power of graphics processing units originally developed for video gaming, and the increasing availability of large datasets.[3] Since 2012, deep learning has brought seismic changes to the technology industry, with major breakthroughs in areas as diverse as computer vision, image caption, speech recognition, natural language translation, robotics, and even self-driving cars.[4–9] In 2015, Scientific American listed deep learning as one of their “world changing” ideas for the year.[10]

Until now, the development and implementation of deep learning methodology into healthcare has faced three main blockers: Firstly, access to large, well-curated, and well-labeled datasets is a requirement that represents a major challenge to the use of deep learning. Although numerous institutions around the world have access to large clinical datasets, far fewer have them in a computationally tractable form and with robust clinical labels for learning tasks. Secondly, highly specialized computing resources are needed, since the performance of deep learning models is highly dependent on recent advances in massively parallel computing architectures, termed graphic processing units. The architecture of silicon customized to these tasks is rapidly evolving with software companies increasingly designing their own hardware chips such as tensor processing units, and field-programmable gate arrays.[11, 12]. Thus, it is already clear that it will be difficult for small research groups, working alone in hospital and university settings, to accommodate these huge financial costs and the rapidly evolving landscape. Thirdly, highly specialized technical expertise and significant mathematical knowledge is required to develop deep learning models. Currently, it is estimated that fewer than 10,000 people worldwide have the skills necessary to tackle serious AI research with the majority of these employed outside academic institutions.[13] According to the 2017 Computer Research Association Taulbee survey, nearly 60% of new AI PhD graduates are recruited by industry.[14]

One approach to combat these obstacles is the increasingly popular technique, transfer learning, where a model developed for a specific task is repurposed and leveraged as a starting point for training on a novel task. While transfer learning mitigates some of the significant computing resources required in designing a bespoke model from inception, it nevertheless requires specialized deep learning expertise to deliver effective results. With this in mind, several companies released application programming interfaces (API) in 2018, claiming to have automated deep learning to such a degree that any individual with basic computer competence could train a high-quality model.[15, 16]

As programming is not a common skill among healthcare professionals, automated deep learning is a potentially promising platform to support the dissemination of deep learning application development in healthcare and medical sciences. In the case of classification tasks, these API products match generic neural-network architectures to a given imaging dataset, fine-tune the network aiming at optimizing discriminative performance, and create a prediction algorithm as output. In other words, the input is a (labeled) image dataset, and the output is a custom classifying algorithm. Yet, the extent to which non-experts can replicate trained deep learning engineers’ performance with the help of automated deep learning remains unclear.

In this study, healthcare professionals without any deep learning expertise explored the feasibility of automated deep learning model development and investigated the performance of these models in diagnosing a diverse range of disease from medical imaging. More precisely, we (i) identified medical benchmark imaging datasets for diagnostic image classification tasks and their corresponding publications on deep learning models; (ii) used these datasets as input; (iii) replaced the classic deep learning models with automated deep learning models; (iv), and compared the discriminative performance of the classic and the automated deep learning models. Moreover, we sought to evaluate the interface that was used for automated deep learning model development (Google Cloud AutoML© Vision API, Beta release) for its utilization in prediction model research. [17–19]

## RESULTS

### Challenge 1: Performance of automated deep learning in archetypal binary classification tasks

Diagnostic properties and discriminative performance were comparable in the case of the investigated binary classification tasks.

#### Task 1: Classification of diabetic retinopathy vs normal retina on fundus images

The Retinal Fundus Image dataset involved 1187 images in total, with 533 normal fundus images (“R0 cases”), and 153 images showing mild (“R1 cases”), 247 moderate (“R2 cases”) and 254 severe diabetic retinopathy (“R3 cases”). Thirteen duplicate images were automatically excluded by the API. The automated deep learning model trained to distinguish healthy fundus images from fundus images showing diabetic retinopathy (R0 from R1, R2 and R3 cases) reached an AUPRC of 0.87, and best accuracy at a cut-off value of 0.5 with a sensitivity of 73.3% and a specificity of 67%.

#### Task 2: Classification of pneumonia vs normal on paediatric CXR

The Paediatric CXR set provided by Guangzhou Medical University/Shiley Eye Institute involved 5827 of 5232 patients CXR images (1582 showing normal pediatric chest x-rays, and 4245 showing pneumonia). The API detected and excluded 8 duplicate images. The AUPRC of this automated deep learning model was 1, best accuracy was reached at a cut-off value of 0.5 with a sensitivity of 97% and a specificity of 100%.

### Challenge 2: Performance of automated deep learning in multiple classification tasks

The two models trained to distinguish multiple classification tasks showed high diagnostic properties and discriminative performance.

#### Task 1: Classification of three common macular diseases and normal retinal OCT images

The Retinal OCT set provided by Guangzhou Medical University/Shiley Eye Institute involved 101418 images of 5761 patients. 31882 images depicted OCT changes related to neovascular age-related macular degeneration (791 patients), 11165 to diabetic macular edema (709), 8061 depicted drusen (713 patients), and 50310 were normal (3548 patients). One hundred seventy fives images were identified as duplicates and excluded by the API. The AUPRC of the automated deep learning model trained to distinguish these four categories was 0.99, while best accuracy was reached at a cut-off value of 0.5, with a sensitivity of 97.3%, a specificity of 100% and a positive predictive value (PPV) of 97.7%.

#### Task 2: Classification of seven distinct categories of skin lesions using dermatoscopic images

The Dermatology Image set involved 10013 images of skin lesions of 10013 patients (327 images depicted actinic keratosis, 514 basal cell carcinoma, 6703 nevus, 1113 melanoma, 115 dermatofibroma, 142 vascular lesion, and 1099 benign keratosis consisting of seborrheic keratosis, solar lentigo and lichen-planus like keratoses). There were no duplicate images detected. The AUPRC of the automated deep learning model trained to distinguish these seven categories was 0.93, while best accuracy was reached at a cut-off value of 0.5, with a sensitivity of 91% and a positive predictive value of 91%.

### Challenge 3: Performance of automated deep learning in a multi-label classification tasks

The automated deep learning model trained to perform a multi-label classification task on the Adult CXR dataset showed poor diagnostic properties and a discriminative performance near chance (AUPRC: 0.57, best accuracy at a cut-off value of 0.5, with a sensitivity of 38% and a positive predictive value of 71%). The NIH CXR14 comprised 11542 cases of atelectasis, 2399 of cardiomegaly, 3323 of consolidation, 1862 of edema, 8036 of effusion, 1734 of emphysema, 1215 of fibrosis, 156 of hernia, 11785 of infiltration, 2923 of mass, 3009 of nodule, 1216 of pleural thickening, 325 of pneumonia, 2199 of pneumothorax and 60304 with no findings of 112120 patients. Twelve duplicates were detected and excluded by the API.

### Comparison with existing state-of-the-art

Subsequently, we compared the diagnostic properties and the diagnostic performance of algorithms trained using automated deep learning on the Retinal Fundus Image, Retinal OCT, Paediatric CXR, Adult CXR, and Dermatology Image datasets compared to best performing deep learning algorithms found in the literature (see Table 2). Interestingly, all best performing algorithms used transfer learning. Some automated deep learning models showed comparable diagnostic properties at a threshold of 0.5 to state-of-the-art deep learning algorithms in published literature. For instance, (i) using the OCT dataset, automated deep learning achieved a sensitivity of 97% and a specificity of 100% (versus a sensitivity of 98% and a specificity of 97% published by Kermany and colleagues); (ii) using the Paediatric CXR dataset, automated deep learning reached a sensitivity of 97% and a specificity of 100% (versus a sensitivity of 93% and a specificity of 90% published by Kermany and colleagues). Other models however, showed lower diagnostic properties, i.e. (i) using the multi-label classification task of the NIH CXR14 dataset in which automated deep learning reached sensitivity of 38%, a positive predictive value of 71% and a AUPRC of 0.57 (versus a AUC of 0.87 published by Guan and colleagues). ; or (ii) using the Retinal Fundus Image dataset, in which automated deep learning reached a sensitivity of 73% and a specificity of 67% (versus a sensitivity of 86% and a specificity of 97% published by Li and colleagues). The thresholds were reported in two cases.[28, 29]

**Table 1.** Summary of the diagnostic properties and the discriminative performance of all five automated deep learning models.

| **MESSIDOR: Fundus images** |  |  |  |  |  |  |  |  |  |
| --- | --- | --- | --- | --- | --- | --- | --- | --- | --- |
| *Classification task* | *Prev* | *TP* | *FP* | *TN* | *FN* | *AUPRC* | *PPV* | *Sens* | *Spec* |
| Presence vs Absence of DR | 55% | 48 | 18 | 36 | 18 | 0.87 | 73% | 73% | 67% |
| **Guangzhou Medical University/Shiley Eye Institute: Retinal OCT images** |  |  |  |  |  |  |  |  |  |
| *Classifications* | *Prev* | *TP* | *FP* | *TN* | *FN* | *AUPRC* | *PPV* | *Sens* | *Spec* |
| Overall | 100% | n.r. | n.r. | n.r. | n.r. | 0.99 | 98% | 97% | 100% |
| CNV vs. Others | 25% | 246 | 2 | 973 | 1 | n.r. | 99% | 100% | 100% |
| Drusen vs. Others | 24% | 208 | 0 | 975 | 23 | n.r. | 100% | 90% | 100% |
| DMO vs. Others | 25% | 247 | 1 | 974 | 0 | n.r. | 100% | 100% | 100% |
| Normal vs. Others | 26% | 250 | 0 | 975 | 0 | n.r. | 100% | 100% | 100% |
| **Guangzhou Medical University/Shiley Eye Institute: Paediatric CXR images** |  |  |  |  |  |  |  |  |  |
| *Classifications* | *Prev* | *TP* | *FP* | *TN* | *FN* | *AUPRC* | *PPV* | *Sens* | *Spec* |
| Pneumonia vs. Normal | 74% | 412 | 6 | 153 | 10 | 1 | 97% | 97% | 100% |
| **NIH CXR14: Adult CXR images** |  |  |  |  |  |  |  |  |  |
| *Classifications* | *Prev* | *TP* | *FP* | *TN* | *FN* | *AUPRC* | *PPV* | *Sens* | *Spec* |
| Overall | n.r. | n.r. | n.r. | n.r. | n.r. | 0.57 | 71% | 38% | n.r. |
| **HAM 10000: Dermatology Image set** |  |  |  |  |  |  |  |  |  |
| *Classifications* | *Prev* | *TP* | *FP* | *TN* | *FN* | *AUPRC* | *PPV* | *Sens* | *Spec* |
| Overall | 100% | n.r. | n.r. | n.r. | n.r. | 0.93 | 91% | 91% | n.r. |
| Actinic keratosis | 3.3% | 25 | 9 | 961 | 8 | n.r. | 74% | 76% | 99% |
| Basal cell carcinoma | 5.2% | 46 | 8 | 943 | 6 | n.r. | 85% | 88% | 99% |
| Nevus | 67% | 656 | 13 | 319 | 15 | n.r. | 98% | 98% | 96% |
| Melanoma | 11% | 75 | 12 | 879 | 37 | n.r. | 86% | 67% | 99% |
| Dermatofibroma | 1.2% | 7 | 3 | 988 | 5 | n.r. | 70% | 58% | 100% |
| Vascular lesion | 1.4% | 13 | 0 | 989 | 1 | n.r. | 100% | 93% | 100% |
| Benign keratosis | 1.1% | 91 | 10 | 883 | 19 | n.r. | 90% | 83% | 99% |

n.r. = not reported
Prev = prevalence: number of given cases as percentage of test dataset
TP = true positives, FP = false positives, TN = true negatives, FN = false negatives
AUPRC = area under the precision-recall curve
PPV = positive predictive value, Sens=sensitivity, Spec = Specificity
DR = Diabetic Retinopathy
CNV = Choroidal neovascularization
DMO = Diabetic macular oedema

**Table 2.** Image classification performance of algorithms trained using automated deep learning compared to best performing algorithms found in the literature.

| **MESSIDOR: Fundus images** |  |  |  |  |
| --- | --- | --- | --- | --- |
|  | *Model architecture* | *Threshold* | *Sensitivity* | *Specificity* |
| Faes & Wagner et al. | Automated deep learning | 0.5 | 73% | 67% |
| Li et al. | VGG-s / Conv1-Fc8 | 0.5 | 86% | 97% |
| **Guangzhou Medical University/Shiley Eye Institute: Retinal OCT images** |  |  |  |  |
|  | *Model architecture* | *Threshold* | *Sensitivity* | *Specificity* |
| Faes & Wagner et al. | Automated deep learning | 0.5 | 97% | 100% |
| Kermany et al. | Inception V3 | n.r. | 98% | 97% |
| **Guangzhou Medical University/Shiley Eye Institute: Paediatric CXR images** |  |  |  |  |
|  | *Model architecture* | *Threshold* | *Sensitivity* | *Specificity* |
| Faes & Wagner et al. | Automated deep learning | 0.5 | 97% | 100% |
| Kermany et al. | Inception V3 | n.r. | 93% | 90% |
| **NIH CXR14: Adult CXR images** |  |  |  |  |
|  | *Model architecture* | *Threshold* | *Sensitivity* | *Specificity* |
| Faes & Wagner et al. | Automated deep learning | 0.5/0.7 | 38%/23% | n.r. |
| Guan et al. | ResNet-50 / DenseNet-121 | 0.7 | n.r. | n.r. |

n.r. = not reported

### Further evaluation of automated model performance

The AutoML©®™ Cloud Vision API provides confusion matrices in the case of single-label classification tasks, in order to uncover label categories in which the model performs insufficiently. The model trained to distinguish the four ophthalmic diagnoses from OCT images (Guangzhou Medical University/Shiley Eye Institute), classified drusen as choroidal neovascularization (CNV) in 10% of cases- implicating a more urgent referral than needed. The model trained on the Dermatology Image set on the other hand, misclassified 28.6% of melanomas as nevus, which in a real-world setting would result in less urgent referral for further work-up and delayed, or worse, missed diagnosis. Moreover, this model also had a high misclassification rate (41.7%) for images showing dermatofibromas. Tables 3 and 4 show the corresponding confusion matrices for the OCT and Dermatology Image set and Fig 1 shows cases from each model where the incorrect label was predicted.

**Fig 1.**
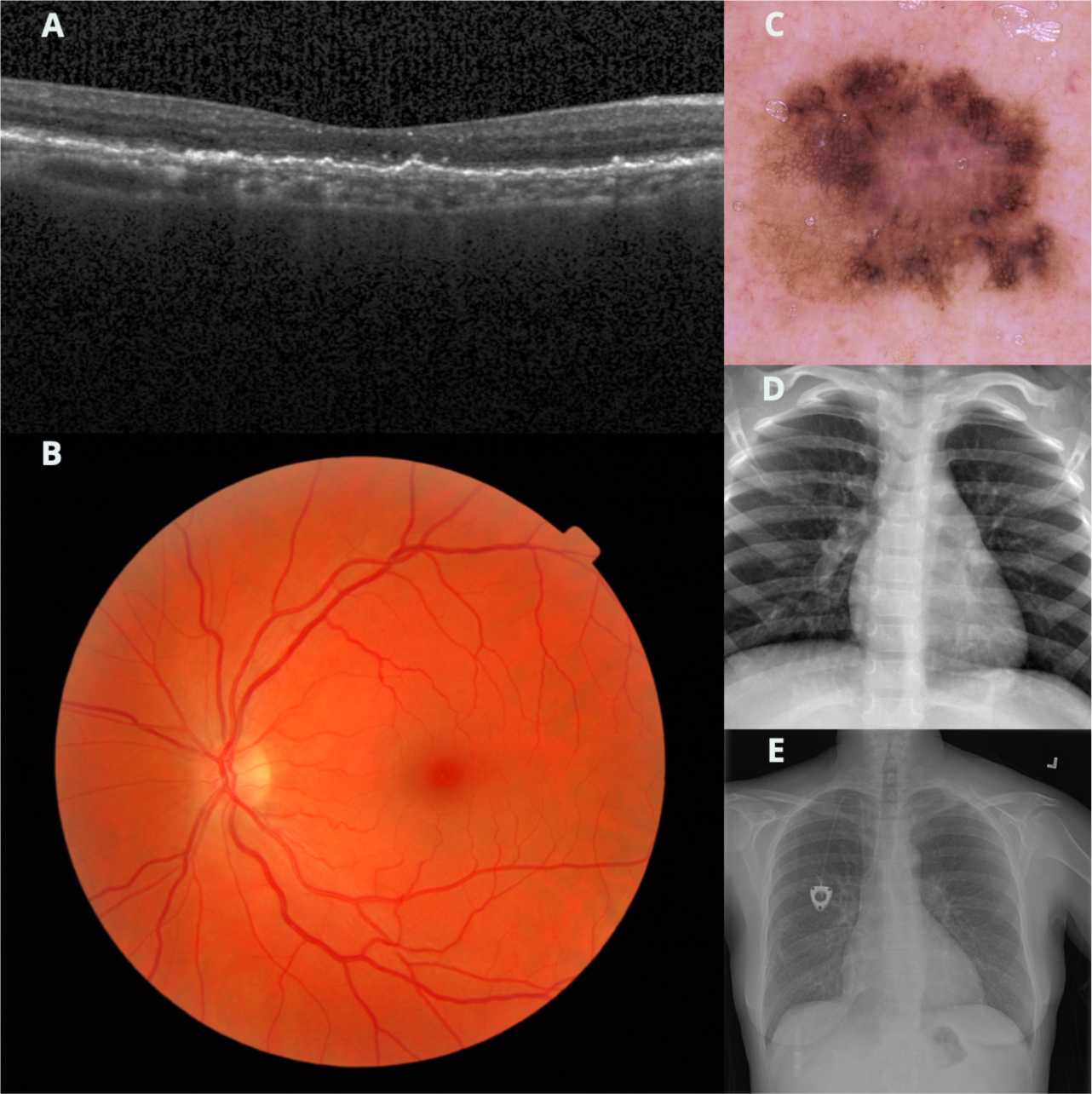
Cases from each model where the incorrect label was predicted. (A) Case of drusen, which was predicted as neovascular age-related macular degeneration. (B) Presence of diabetic retinopathy predicted as normal. (C) A melanoma predicted as a nevus. (D) Pneumonia predicted as normal. (E) A pleural effusion predicted as normal. Case B does not have detectable features of its label while E is equivocal.

**Table 3.** The confusion matrix for the model developed on the OCT image dataset provided by Guangzhou Medical University/Shiley Eye Institute.

|  |  | Predicted label |  |  |  |
| --- | --- | --- | --- | --- | --- |
|  |  | *Drusen* | *CNV* | *DMO* | *Normal* |
| True label | *Drusen* | 0.9 | 0.1 | - | - |
|  | *CNV* | - | 0.996 | 0.004 | - |
|  | *DMO* | - | - | 1 | - |
|  | *Normal* | - | - | - | 1 |

CNV = Choroidal neovascularization
DMO = Diabetic macular oedema

**Table 4.** The confusion matrix for the model developed on the Dermatology Image set.

|  |  | Predicted label |  |  |  |  |  |  |
| --- | --- | --- | --- | --- | --- | --- | --- | --- |
|  |  | *Melanoma* | *Naevus* | *Benign keratosis* | *Actinic keratosis* | *BCC* | *Dermato-fibroma* | *Vascular skin lesion* |
| True label | *Melanoma* | 0.67 | 0.286 | 0.027 | - | 0.018 | - | - |
|  | *Naevus* | 1 | 0.978 | 0.003 | 0.001 | 0.004 | 0.003 | - |
|  | *Benign keratosis* | 0.027 | 0.091 | 0.827 | 0.045 | - | 0.009 | - |
|  | *Actinic keratosis* | - | 0.061 | 0.121 | 0.758 | 0.061 | - | - |
|  | *BCC* | 0.019 | 0.019 | 0.038 | 0.038 | 0.885 | - | - |
|  | *Dermatofibroma* | 0.167 | 0.167 | - | 0.083 | - | 0.583 | - |
|  | *Vascular skin lesion* | - | - | - | - | 0.071 | - | 0.929 |

### External Validation

In the case of the deep learning model developed on a subset of the Dermatology Image set, we additionally performed an external validation using the Dermatology Validation set. As the latter set did not include benign keratosis as a label, these images were removed from the Dermatology Image set used for training.

The automated deep learning model showed poor diagnostic properties and a discriminative performance near chance (AUPRC: 0.47, best accuracy at a cut-off value of 0.5, with a sensitivity of 49% and a positive predictive value of 52%). Of note, the sensitivity for melanoma classification is 11% with a misclassification rate of 63.7%.

Interestingly, nevus was the most likely classification in all cases, followed by the ground truth. The only exception was the case of actinic keratosis, where its ground truth diagnosis was the third most probably diagnosis to be predicted (10.6% likelihood). The prevalence of images showing nevus was 76% in the developmental dataset, compared to 36% in the dataset used for external validation. To investigate the impact of class imbalance between the Dermatology Image set and Dermatology Validation Set, we undersampled the nevus class by 3000 images and assessed the resulting change in accuracy. Only minimal improvements were noted in its discriminative performance and diagnostic properties (data available upon request).

Tables 5a and 5b show the external validation of the model trained on a subset of the Dermatology Image set; (a) discriminative performance and diagnostic properties; (b) confusion matrix.

**Table 5a.**
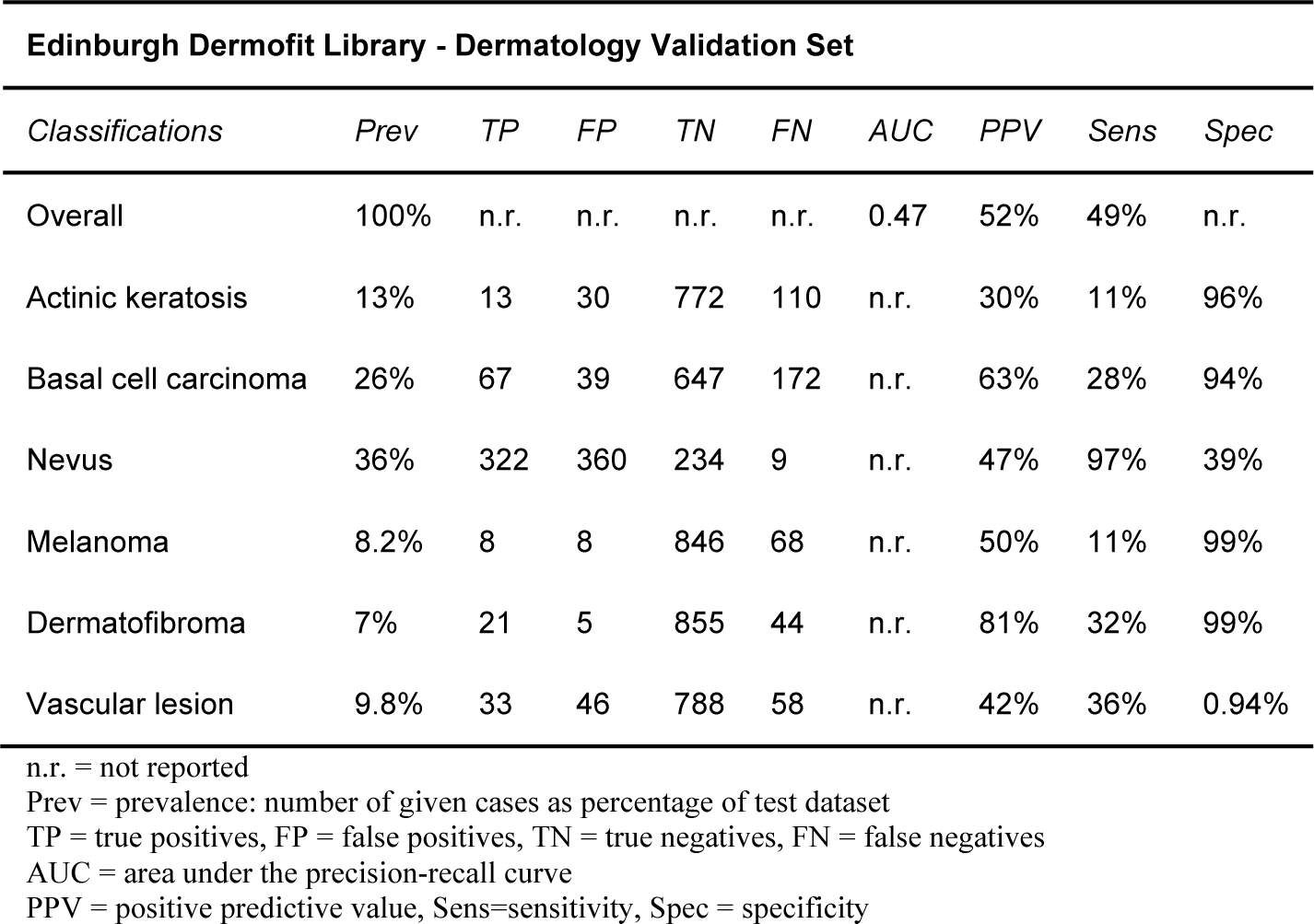
The diagnostic properties and discriminative performance of the external validation of the algorithms trained using automated deep learning on the Dermatology Image set.

**Table 5b.** The confusion matrix of the external validation of HAM10000-trained algorithm on the Edinburgh Dermofit Library dataset.

|  |  | Predicted label |  |  |  |  |  |
| --- | --- | --- | --- | --- | --- | --- | --- |
|  |  | *BCC* | *Naevus* | *Actinic keratosis* | *Melanoma* | *Vascular skin lesion* | *Dermatofibroma* |
| **True label** | *BCC* | 0.28 | 0.552 | 0.113 | 0.008 | 0.042 | 0.004 |
|  | *Naevus* | 0.003 | 0.973 | 0.003 | - | 0.009 | 0.012 |
|  | *Actinic keratosis* | 0.203 | 0.683 | 0.106 | 0.008 | - | - |
|  | *Melanoma* | 0.039 | 0.789 | 0.013 | 0.118 | 0.039 | - |
|  | *Vascular skin lesion* | 0.033 | 0.582 | - | 0.022 | 0.363 | - |
|  | *Dermatofibroma* | 0.108 | 0.477 | 0.015 | 0.031 | 0.046 | 0.323 |

BCC=Basal cell carcinoma

## DISCUSSION

### Main findings

This manuscript reports a first of its kind implementation of automated deep learning, in which we demonstrate that medical professionals with limited programming experience can utilize automated deep learning to develop algorithms that can perform clinical classification tasks to a level comparable with traditional deep learning models that have been applied in existing literature. Most of the automatically developed deep learning models, except for that trained on the multi-label classification task of the Adult CXR set, showed comparable discriminative performance and diagnostic properties to state-of-the-art performing deep learning algorithms.

From a methodological viewpoint, our results – as with those reported in the state-of-the-art deep learning studies – might be overly optimistic, since we were not able to test all the models “out-of-sample” as recommended by current guidelines. Moreover, for external validation, the present version of the API only allows single image upload for prediction, limiting large scale external validation. This reduces its usability for systematic evaluation in prediction model research considerably, given the high numbers of images that these datasets comprise. To circumvent this issue, we created a proxy to an external validation and found a substantial reduction in the diagnostic and discriminative performance of the deep learning model. The limited performance of automated deep learning models (also in the multi-label classification task) are likely related to inadequacy of the datasets used to train the models. To obviate concerns over class imbalance in our external validation, we under-sampled accordingly, however, this did not significantly alter the model’s discriminative performance.

### Strengths

To our knowledge, this is the first assessment of the feasibility and usefulness of automated deep learning technology in medical imaging classification tasks performed by healthcare professionals. In this study, we showed best effort to comply with the reporting guidelines for prediction model research, and for developing and reporting machine learning predictive models. [24, 30] One major strength of our study is that we tested one exemplary model for robustness in an out-of-sample cross validation, since internal validation and random split sample-validation has been claimed to overestimate test performance. A further strength is that our results can easily be explored and replicated by others, given the use of public datasets and the free trial use of the the AutoML©®™ Cloud Vision API.

### Limitations and future directions

The convenience sampling used in the out-of-sample cross validation has been claimed to introduce bias and exaggerate estimates of diagnostic performance. [31] Furthermore, the API was not able to depict saliency maps and consequently we were not able to interrogate the model for the image areas which it considered most important for its prediction.[32] This “black-box” classification does not provide any biological information useful for clinical purposes.[33] Moreover, the specifics of the models used by the API are not transparent, which constitutes a barrier to their evaluation and the reproducibility of this study.[34] We were not able to extract or calculate all metrics and measures of uncertainty conventionally used in prediction model research in all cases (i.e. specificity or confidence intervals), which impedes comparison to the current best technology other than deep learning. We noted that best accuracy was consistently achieved at a threshold of 0.5 by the API. This is likely the default threshold to which the API is optimized: a setting which is inaccessible to the user during model development. In clinical practice, thresholds should be set according to the role of the diagnostic test and the consequences of a misdiagnosis. Therefore, the ability to adjust the preferred threshold is an important function for creating a fit-for-purpose API.

Besides Google’s AutoML©®™ Cloud Vision API, a number of vendors recently have released similar automated deep learning platforms including both established technology corporations (e.g. Amazon SageMaker, Baidu EZDL, IBM Watson Studio) and start-ups (Oneclick.ai, Platform.ai). Our study pertained to only one API, Google’s AutoML©®™, as this was among the first open neural architecture search-based engines released and was freely available on a trial basis. While this report is a proof-of-concept evaluation of healthcare professional-led deep learning, it is unclear whether other APIs may provide greater discriminative performance. Assessment of other platforms is an objective of our future research.

Currently, studies on AutoML including ours, have to rely on publicly available datasets. While using them allows comparison of performance between different algorithms, these are not without concern. For many classification tasks, and particularly for validation purposes, the existing datasets tend to be too small and not representative of the general population. Moreover, data quality in general could be improved. A full evaluation of dataset limitations is beyond the scope of this manuscript, however inconsistent labeling and duplicate images bore direct pertinence to our study. Issue such as equivocal labels (figure), image confounders (presence of chest drain in images of pneumothorax), and label semantics (nodule vs mass, consolidation vs pneumonia) have been noted previously in datasets used for deep learning.[35] Apart from the Dermatology Image set, all datasets contained duplicate images. Conveniently, Google’s AutoML©®™ Cloud Vision API will automatically detect, indicate and exclude the relevant images, however clinicians need to be cognizant to this possibility as other APIs may lack this feature and generate spurious evaluation metrics. Since the quality of the results obtained using deep learning models substantially depends on the quality of the dataset employed in the model development, it would therefore be imperative that patient demographics and information about the way the data was collected (i.e. patient flow) would be presented as the validity and generalization of the models cannot otherwise be assessed. In our study, we were only able to provide limited information about the descriptives of these datasets, using what has been published by their creators.

With the availability of new and carefully administered datasets, many validity problems could be resolved. Great hopes lay in data sharing initiatives as promoted from many peer-reviewed journals or those from the Dataverse Network Project and the UK Data Archive. [36, 37] On the other hand, these initiatives struggle with issues of confidentiality and anonymity when publishing or sharing data relating to individuals. Moreover, regulatory restrictions still remain. Fortunately, recent developments both in the United Kingdom with the NHS Digital Guidance and the call for Health Insurance Portability and Accountability Act compliance in the United States have clarified the framework for many public Cloud systems.[38, 39] The European Union General Data Protection Regulation is another possible barrier towards an efficient use of published data. However, since many studies will be dealing with ephemeral processing of de-identified data we do not believe that the General Data Protection Regulation is likely to pose a substantial hindrance.

We confirmed feasibility but encountered various methodological problems that are well-known in research projects performing classification tasks and predictions. We believe that concerted efforts in terms of data quality and accessibility are needed to make automated deep learning modelling successful. Moreover, as the technology evolves, transparency in the reporting of the models and a careful reporting of their performance in methodologically sound validation studies will be pivotal for a successful implementation into practice. Finally, the extent to which automated deep learning algorithms must adhere to regulatory requirements for medical devices is unclear. [34]

Although the development of deep learning prediction models was feasible for healthcare professionals without any deep learning expertise, we would recommend the following developments for automated deep learning: (i) transparency of the model architectures and technical specifications in use; (ii) reporting of established performance parameters in prediction model research, such as sensitivity, specificity (iii) reporting of the label distribution in automatically conducted random-split sample validations (in cases, in which the subsets have not been stipulated by the user explicitly); (iv) depiction of all incorrectly and correctly classified images (true positive, false negative, false positive, true negative cases); and (v) a robust solution to allow systematic external validation.

### Conclusion

The availability of automated deep learning may be a cornerstone for the democratization of sophisticated algorithmic modelling in healthcare as it allows the derivation of classification models without requiring a deep understanding of the mathematical, statistical and programming principles. However, the translation of this technological success to meaningful clinical impact constitutes requires concerted efforts and a careful stepwise work-up. The sharp contrast of the model’s discriminative performance in internal versus external validation may foretell the ultimate use case for automated deep learning software once the technology matures. As researchers and clinicians have excellent access to images and patient data within their own institutions, they may be able to design auto ML models for internal research, triage, and customized care delivery. This may avert the need for costly external prospective validation across imaging devices, patient populations and clinician practice patterns. In contrast, large scale standardized deep learning algorithms will necessitate worldwide, multi-variable validations of expertly tuned models. Thus, there is considerable value to these “small data” approaches customized to a specific geographical patient population that a given clinic may encounter. This may be where automated deep learning finds its niche in the medical field. Importantly, this could make models susceptible to selection bias, over-fitting, and a number of other issues from imprecise model training or patient selection. Therefore, regulatory guidelines are needed for both medical deep learning and clinical implementation of these models before they may be used in clinical practice. In summary, while our approach seems rational in this early evaluation, the results of this study cannot yet be extrapolated into clinical practice.

## METHODS

### Study design and data source

We used five distinct open-source datasets comprising medical imaging material to automatically develop five corresponding deep learning models for the diagnosis of common diseases or disease features. Namely, we trained deep learning models on; (1) retinal fundus images (the MESSIDOR dataset, hereafter referred to as Retinal Fundus Image set); (2) retinal optical coherence tomography (OCT) images (Guangzhou Medical University/Shiley Eye Institute Version 3, hereafter referred to as Retinal OCT set); (3) pediatric chest X-ray (CXR) images (Guangzhou Medical University/Shiley Eye Institute, hereafter referred to as Paediatric CXR set); (4) adult CXR images (the National Institute of Health (NIH) CXR14 dataset, hereafter referred to as Adult CXR set), and; (5) dermatology images (the Human against Machine (HAM) 10000 dataset, hereafter referred to as Dermatology Image set).[20–23] Moreover, in a proof-of-principle evaluation, we aimed at testing one of the models “out-of-sample” as recommended by current guidelines.[24] The current version of the API only allows single image upload for model prediction limiting the feasibility of large scale external validation. However, we created a proxy to an external validation in one exemplary use-case, where we used the Dermatology Image set for algorithm development and tested its performance using the Edinburgh Dermofit Library dataset (hereafter referred to as Dermatology Validation set).[25]

### Training of healthcare professionals using the API

Two clinicians (LF and SKW) with no prior coding or machine learning experience performed the analysis after a period of self-study based on the online documentation. In total, both researchers invested approximately ten hours of preparation. Due to the release cycle evolution of the AutoML©®™ Cloud Vision API during the study (alpha release May 2018, beta release July 2018), they adopted an iterative approach when executing the analyses. All analytic steps and interpretations of results were performed jointly.

### Patient recruitment and enrolment

We accessed five de-identified open-source imaging datasets that were collected from retrospective, non-consecutive cohorts, showing diseases or disease features of common medical diagnoses. Eligibility criteria, patients’ demographics and patient workflow for each of these datasets are published elsewhere.[20–23]

### Index Test: AutoML©®™ Cloud Vision API

The term “automated machine learning” commonly refers to “automated methods for model selection and/or hyperparameter optimization”. This is the concept which led to the idea of allowing a neural network to design another neural network, through the application of a neural architecture search.[17–19] In deep learning, designing and choosing the most suitable model architecture requires a significant amount of time and experimentation even for those with significant deep learning expertise. This is because the search space of all possible model architectures can be exponentially large (e.g. a typical 10-layer network could have ∼10^10^ candidate networks). To make this model design process easier and more accessible, an approach known as neural architecture search has recently been described. [26] Neural architecture search is typically achieved using one of two methods: 1) reinforcement learning algorithms, and 2) evolutionary algorithms. The former forms the basis of this commercially available API that was evaluated in this study.[27]

### Data handling and analytic approach

We uploaded images of five open-source datasets to a Google Cloud bucket in conjunction with a comma-separated value file indicating the image label, file-path and dataset distribution (i.e. training-, validation- or test dataset). Images were allocated to the training, validation and test datasets (80%, 10% and 10% respectively) using a random number function. In the case of the Retinal OCT images where a specific test set had been stipulated in a previous report, we compared our performance to that published by using the same test set.[21] Duplicate images were automatically detected and excluded by the API. We did not relabel any of the used datasets. All models were trained for a maximum of 24 compute hours. Except for the Retinal OCT set, the discriminative performance of each deep learning model was evaluated using the randomly specified test dataset, and in the case of the deep learning model developed on a subset of the Dermatology Image set, additionally in an external validation using an independent open-source dermatology dataset (Edinburgh Dermofit Library).

### Comparison with Benchmark Classic Deep Learning Models

In order to provide a direct comparator to the performance of classic deep learning models developed using traditional non-autoML techniques (deep learning models with bespoke architectures for a data and problem set developed by human experts), we conducted a systematic search of the literature to identify classical deep learning models, composed by deep learning experts, which have been trained and/or validated on the five open-source datasets. The performance of these existing models served as a direct comparator with the API. We searched (Pre-)Medline, Embase, Science Citation Index, Conference Proceedings Citation Index, Google Scholar and arXiv document server from 01 January 2012 until 05 October 2018. Studies were included if they developed a deep learning algorithm on the datasets used in this study. No language restrictions were applied. The search strategy is available as a supplementary attachment (see Supplementary File 1). We pre-specified the cut-off of 2012 on the basis of a step-change in deep learning performance; a deep learning model called AlexNet, won a visual recognition challenge, the ImageNet Large-Scale Visual Recognition Challenge, for the first time.[4] If a study provided contingency tables for the same or for separate algorithms tested in a specific classification task, we assumed these to be independent from each other. We accepted this, as we were particularly interested in providing an overview of the results of various studies rather than providing accurate precisions of point estimates.

### Statistical Analysis

The AutoML©®™ Cloud Vision API provides metrics that are commonly used by the AI-community. These are recall (sensitivity) or precision (positive predictive value) for given thresholds and the area under the precision recall curve (AUPRC). Additionally, confusion matrices are depicted for each model, cross-tabulating ground truth labels versus the labels predicted by the deep learning model. Where possible, we extracted binary diagnostic accuracy data and constructed contingency tables and calculated specificity at the threshold of 0.5. Contingency tables consisted of true-positive (TP), false-positive (FP), true-negative (TN) and false-negative (FN) results. For consistency, we adhered to the typical test accuracy terminology: sensitivity (recall), specificity and positive predictive value (precision). The classification tasks were chosen according to their popularity in the current AI literature for the purpose of comparability to state-of-the-art deep learning models. Where possible, we plotted contingency tables against the ones reported by other studies using the same benchmark datasets to develop deep learning models.

We *a priori* attempted to compare the classification performance between state-of-the-art deep learning studies and our results. However, while the published reports provided areas under the receiver operating characteristic curve (AUC ROC), the AutoML API reports the AUPRC. Although the points of the two types of curves can be mapped one-to-one and hence curves can be translated from the ROC space to the prediction space (if the confusion matrices are identical) differences in the confusion matrices and the level of reporting impeded us from performing a comparison on the level of AUC. Instead, we compared the performance on the level of sensitivity and specificity at the same threshold as had been used in the previous reports.

## AUTHOR CONTRIBUTIONS

All authors contributed in the conception and design of the study. LF and SW trained the automated deep learning models and collected the data, which was analyzed by LF, SW, DF, LB and PK. LF and SW drafted the manuscript, which was revised with significant input from all authors. All authors have approved the final version of the article.

